# Fussing about fission: defining variety among mainstream and exotic apicomplexan cell division modes

**DOI:** 10.1101/2020.04.23.056333

**Authors:** Marc-Jan Gubbels, Caroline D. Keroack, Sriveny Dangoudoubiyam, Hanna L. Worliczek, Aditya S. Paul, Ciara Bauwens, Brendan Elsworth, Klemens Engelberg, Daniel K. Howe, Isabelle Coppens, Manoj T. Duraisingh

## Abstract

Cellular reproduction defines life, yet our textbook-level understanding of cell division is limited to a small number of model organisms centered around humans. The horizon on cell division variants is expanded here by advancing insights on the fascinating cell division modes found in the Apicomplexa, a key group of protozoan parasites. The Apicomplexa display remarkable variation in offspring number, whether karyokinesis follows each S/M-phase or not, and whether daughter cells bud in the cytoplasm or bud from the cortex. We find that the terminology used to describe the various manifestations of asexual apicomplexan cell division emphasizes either the number of offspring or site of budding, which are not directly comparable features and has led to confusion in the literature. Division modes have been primarily studied in two human pathogenic Apicomplexa, malaria-causing *Plasmodium* spp. and *Toxoplasma gondii*, a major cause of opportunistic infections. *Plasmodium* spp. divide asexually by schizogony, producing multiple daughters per division round through a cortical budding process, though at several life-cycle nuclear amplifications are not followed by karyokinesis. *T. gondii* divides by endodyogeny producing two internally budding daughters per division round. Here we add to this diversity in replication mechanisms by considering the cattle parasite *Babesia bigemina* and the pig parasite *Cystoisospora suis. B. bigemina* produces two daughters per division round by a ‘binary fission’ mechanism whereas *C. suis* produces daughters through both endodyogeny and multiple internal budding known as endopolygeny. In addition, we provide new data from the causative agent of equine protozoal myeloencephalitis (EPM), *Sarcocystis neurona*, which also undergoes endopolygeny but differs from *C. suis* by maintaining a single multiploid nucleus. Overall, we operationally define two principally different division modes: internal budding found in cyst-forming Coccidia (comprising endodyogeny and two forms of endopolygeny) and external budding found in the other parasites studied (comprising the two forms of schizogony, binary fission and multiple fission). Progressive insights into the principles defining the molecular and cellular requirements for internal versus external budding, as well as variations encountered in sexual stages are discussed. The evolutionary pressures and mechanisms underlying apicomplexan cell division diversification carries relevance across Eukaryota.

**Contribution to the Field:** Mechanisms of cell division vary dramatically across the Tree of Life, but the mechanistic basis has only been mapped for several model organisms. Here we present cell division strategies across Apicomplexa, a group of obligate intracellular parasites with significant impact on humans and domesticated animals. Asexual apicomplexan cell division is organized around assembly of daughter buds, but division forms differ in the cellular site of budding, number of offspring per division round, whether each S-phase follows karyokinesis and if mitotic rounds progress synchronously. This varies not just between parasites, but also between different life-cycle stages of a given species. We discuss the historical context of terminology describing division modes, which has led to confusion on how different modes relate to each other. Innovations in cell culture and genetics together with light microscopy advances have opened up cell biological studies that can shed light on this puzzle. We present new data for three division modes barely studied before. Together with existing data, we show how division modes are organized along phylogenetic lines and differentiate along external and internal budding mechanisms. We also discuss new insights into how the variations in division mode are regulated at the molecular level, and possess unique cell biological requirements.

## 1 Introduction

Reproduction is critical for perpetuating a species and lies at the core of the definition of life. Yet, the modes by which cell division can occur are very diverse. The molecular and cellular mechanisms underlying such differences have been dissected for a limited set of model organisms, most of which carry resemblance to mammalian/human cell division. The Apicomplexa comprise a protozoan phylum harboring human pathogens, like malaria-causing *Plasmodium* spp., opportunistic parasites like *Toxoplasma gondii* and *Cryptosporidium* spp., and emerging pathogens like *Babesia* spp. (phylogeny in Fig 1). In addition, many are of economic relevance in agriculture and companion animals such as *Babesia* spp, *Theileria* spp. and *Eimeria* spp. Furthermore, many Apicomplexa infect other birds, mammals, reptiles, amphibians, fish, and invertebrates, but their cell biology has only been studied minimally. The asexual multiplication cycles of the Apicomplexans are diverse, and their sexual cycles add even more diversity. The well-studied Apicomplexa relevant to humans display a diverse spectrum of asexual cell division strategies both within and among species, providing an excellent opportunity for comparative biology. Indeed, we posit that by studying the principles in more distant organisms such as protozoa, which span a wide breadth of evolutionary history and correspondingly diverse biology, we can obtain fascinating new insights and principles.

**Figure 1.**
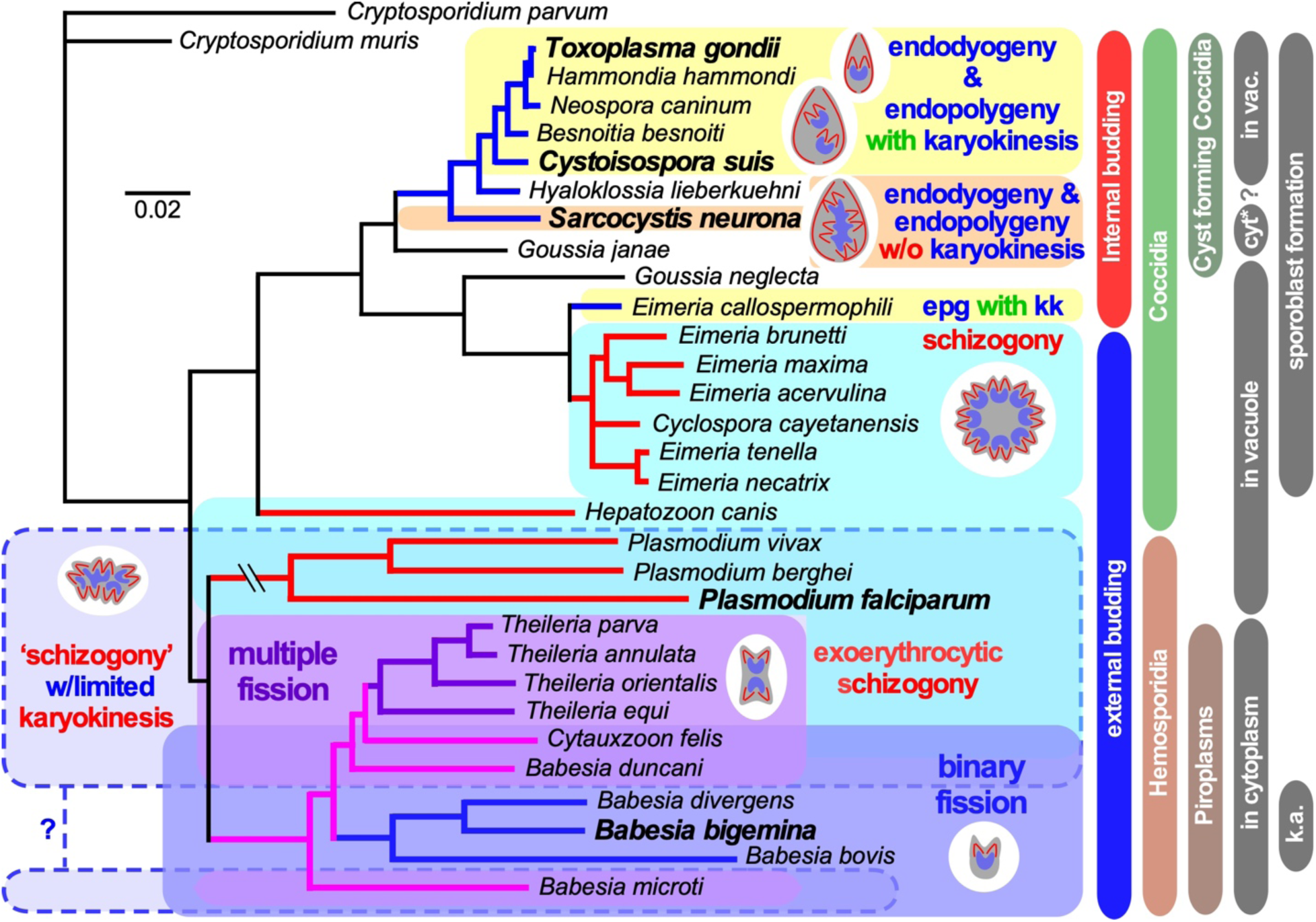
Select apicomplexan phylogeny and division modes. 18S ribosomal RNA based phylogeny of species whose division modes have been studied. *Cryptosporidium* spp. were used as outgroup. Bars on the right indicate different naming and biological relationships, with asexual division modes in blue and red. “in vacuole” and “in cytoplasm” indicate whether asexual replication occurs in a parasitophorous vacuole, or whether the parasite escapes from the vacuole and resides in the cytoplasm of the host cell for its replication. Note that only the acute stage merozoites of *S. neurona* replicates by endopolygeny in the cytoplasm, whereas metrocytes preceding the bradyzoites as well as the bradyzoites divide by endodyogeny within a vacuole supporting a proteoglycan cyst wall. Furthermore, tissue cysts for *C. suis* have not been described and this lacking ability is likely a secondary loss. In addition, for several *Plasmodium* spp. (Simonetti, 1996), as well as *Babesia* and *Theileria* spp. (Jalovecka et al., 2018) it has been shown that sporozoite formation progresses without karyokinesis to produce large polyploid nuclei while budding is from the cortex. We note that *Plasmodium* sporozoites infect hepatocytes wherein they divide by schizogony and manipulate the hepatocytes to expand their size, whereas *Theileria* sporozoites infect white blood cells, replicate by schizogony (Shaw and Tilney, 1992) and trigger white blood expansion as well as division (i.e. transformation, which basically is leukemia (Luder et al., 2009; Chakraborty et al., 2017), which contrasts with *Babesia* sporozoites as they directly infect red blood cells. “epg with kk” means “endopolygeny with karyokinesis”, in case of *E. callospermophili*;”k.a.” means “kinete amplification”, in case of select *Babesia* spp.

Asexual apicomplexan cell division revolves around variations in budding through the assembly of a membrane skeleton that ultimately underlies the plasma membrane. The beginning- and end-point of a cell division round across all Apicomplexa is a “zoite”, a cell type with the phylum-defining complex of apical secretory organelles and cytoskeletal structures (Leander and Keeling, 2003) (e.g. Fig 2A, 3F1, 4A, 5E1, 6C1). These apical structures uniformly facilitate host-cell invasion, an essential step in the obligate intracellular life style of apicomplexan parasites (Gubbels and Duraisingh, 2012; Sharma and Chitnis, 2013; Frenal et al., 2017a). The cortical cytoskeleton just below the plasma membrane, however, plays an additional role as a key structure facilitating budding across asexual division modes for Apicomplexa (Anderson-White et al., 2012; Kono et al., 2016). From the outside in, it is built up from three uniformly conserved elements: alveolar vesicles (alveoli) that anchor the myosin motor enabling motility (Frenal et al., 2017a), an epiplastin (a.k.a. alveolar) protein meshwork (Goodenough et al., 2018), and a series of sub-pellicular, longitudinal microtubules emanating from the apical end. The number and length of the microtubules vary across parasite species and stage (Spreng et al., 2019), but the alveoli and epiplastin meshwork are universally conserved and make up the inner membrane complex (IMC) (Kono et al., 2012). The epiplastin family proteins contain Val-Pro-Val (VPV) repeats and are generally known as alveolins or IMC proteins (Gould et al., 2008; Anderson-White et al., 2011; Kono et al., 2012; Al-Khattaf et al., 2015). Cytoskeletal assembly initiates from the centrosome and all three elements are simultaneously assembled in an apical to basal direction (Chen and Gubbels, 2013; Francia and Striepen, 2014; Suvorova et al., 2015). Another important and shared feature is that across asexual division modes, budding appears to be tied always to a round of S-phase and mitosis (Francia and Striepen, 2014; Suvorova et al., 2015). A fourth cytoskeleton element that is conserved across division modes is the basal complex, a ring residing at the basal end of the daughter bud (Ferguson et al., 2008). This ring contains MORN1, a scaffolding protein that is essential for maintenance of the daughter bud’s basal end integrity (Gubbels et al., 2006; Hu, 2008; Heaslip et al., 2010; Lorestani et al., 2010; Kono et al., 2016). Proteins with repetitive MORN domains are often found in cilia and flagella (Mecklenburg, 2007; Shetty et al., 2007; Tokuhiro et al., 2008; Morriswood and Schmidt, 2015); indeed all apicomplexan division modes share several features with cilia or flagella assembly as seen across eukaryotes. For example, striated fiber assembly (SFA) fibers are found in flagellar assembly in algae to orient the basal body in the flagellum, which in Apicomplexa anchor the centrosome in the daughter bud (Francia et al., 2012). A spindle assembly abnormal protein 6 (SAS6)-like protein found in flagellar basal bodies is also found in the apical polar ring (APR), the microtubule organizing center (MTOC) nucleating the subpellicular microtubules (and conoid, if present) (de Leon et al., 2013; Francia et al., 2015; Wall et al., 2016). Finally, the base of cilia and flagella is often decorated with a contractile centrin and a myosin (Roberts et al., 2004; Trojan et al., 2008), which, at least in *T. gondii*, are represented by TgCentrin2 and Myosin I and J (Hu et al., 2006; Hu, 2008; Frenal et al., 2017b).

**Figure 2.**
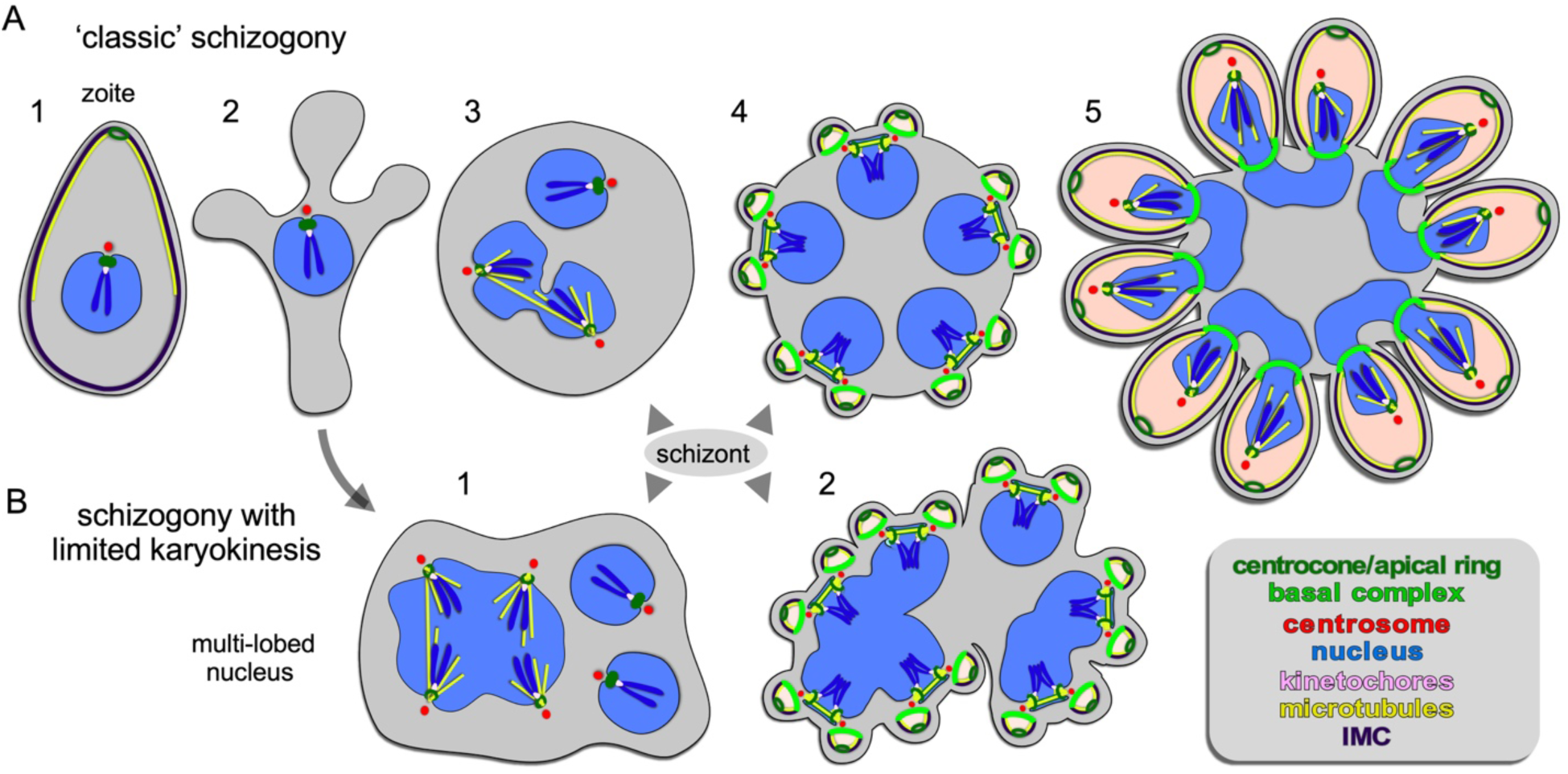
Apicomplexan asexual cell division by schizogony. **A**. Schematic representation of progressive cell division steps during ‘classic’ schizogony. The mother’s cytoskeleton (present in 1) is disassembled following successful host cell invasion resulting in a pleomorphic cell (2) before onset of mitosis and karyokinesis, which can be asynchronous (3). Daughter cells bud from the cortex (4/5), positioned by the centrosome anchored at the plasma membrane (4). **B**. Schematic representation of phases in the cell division during schizogony with limited karyokinesis as seen during Heamosporidian sprorogenesis. Karyokinesis is inconsistently following each S/M-phase leading to a nuclei population with varying levels of ploidity. Note that nuclear cycles within the same nucleoplasm are similar (1), whereas budding is synchronous for all nuclei and linked to a final round of S/M (2). Note that the number of offspring in both forms of schizogony can reach into the 1000s, which is not represented in the schematics. Chromosome condensation does not occur and the chromosomes are only drawn to convey the principle of spindle pole attachment. Mother cytoplasm represented in grey, daughter cytoplasms in pink.

Despite this central plan, many variations exist on what precedes daughter budding, and how daughter bud formation is orchestrated. The first variable is the number of offspring per mother cell in an asexual apicomplexan division round, which can be as few as two or nearly 100,000. The principle behind the uncoupling of daughter formation from S/M-phase was recently established in *T. gondii* by the presence of a bipartite centrosome: the centrosome’s inner-core proximal to the nucleus regulates the S/M cycle independently of the centrosome’s outer-core distal from the nucleus, which organizes daughter bud formation (Chen and Gubbels, 2013; Suvorova et al., 2015; Chen and Gubbels, 2019). Moreover, cyclins and cyclin-dependent kinases (CDKs), as found in higher eukaryotes, ultimately control cell cycle progression but their nature and modes of operation are specifically tailored to the needs of different apicomplexan cell cycle variation (e.g. (Roques et al., 2015; Alvarez and Suvorova, 2017; Ganter et al., 2017; Naumov et al., 2017), recently reviewed in (Matthews et al., 2018; White and Suvorova, 2018)). A second variable is whether karyokinesis follows each S-phase and mitosis (S/M-phase) or not. Finally, daughters can either assemble within the mother’s cytoplasm (internal budding), or assemble and emerge directly from the plasma membrane of the mother cell (cortical or peripheral budding). Strikingly, these modes can vary not only between Apicomplexa, but also across different life-cycle stages within a single species.

Although these insights describe the key principles, the details on cell biological mechanisms facilitating the various division modes are intermittent and often compounded by historical terminology which either do not accurately capture the shared principles or clearly define differences. Here we highlight the best-understood cell division modes, schizogony in *Plasmodium* spp. and endodyogeny in *T. gondii*, supplemented with new data on emerging systems displaying different forms of endopolygeny in *Cystoisopora suis* and *Sarcocystis neurona*, as well as binary fission in *Babesia bigemina*. Emerging insights are subsequently used to assess their kinship and to chart principles underlying these distinct cell division modes.

## 2 Material and Methods

### 2.1 *Babesia* spp. culture

*B. bigemina* strain JG-29 (Mexico) was kindly provided by Dr. David Allred (University of Florida) and grown as described previously (Vega et al., 1985) with modifications as described here. Parasites were culture adapted over a month to grow efficiently in tissue culture media RPMI-1640 supplemented with 25 mM HEPES, 50 mg/l hypoxanthine, 2.42 mM sodium bicarbonate, and 4.31 mg/ml AlbuMAX-II (Invitrogen). Before addition of AlbuMAX-II and sodium bicarbonate, the pH was adjusted to 6.75. Parasites were grown under microaerophilous stationary phase culture conditions with a hypoxic atmosphere of 1:5:94 oxygen: carbon dioxide: nitrogen at 4% hematocrit in washed, defibrinated bovine red blood cells (Hemostat Labs, Dixon, CA).

*B. divergens* strain Rouen 1987, a kind gift from Drs. Kirk Deitsch and Laura Kirkman (Weill Cornell Medical College), was grown in human erythrocytes as described (Paul et al., 2016). Human erythrocytes were purchased from Research Blood Components (Boston, USA).

### 2.2 *Cystoisospora suis* culture

Oocysts of *C. suis* strain Wien-I (Austria) were isolated from porcine fecal samples and used for *in vitro* culture in intestinal porcine epithelial cells (IPEC-J2, ACC 701, Leibniz Institute DSMZ GmbH, Braunschweig, Germany) as described previously (Worliczek et al., 2013) with modifications as described here. For the production of glass cover slips with confluent layers of host cells infected with *C. suis*, cover slips were placed into 6-well culture plates for suspension cultures (PAA, Pasching, Austria). IPEC-J2 were suspended in DMEM/Ham’s F12 medium (PAA, Austria) supplemented with 5 % fetal calf serum, 2 mM L-glutamine, 100 U/ml penicillin and 0.1 mg/ml streptomycin, and seeded in a density of 4×10^5^ cells/well. After overnight incubation at 37°C, 5 % CO_2_ cover slips with attached IPEC-J2 were transferred to 6-well surface-treated culture plates for adherent cells (PAA, Austria). Sporozoites of *C. suis* were excysted as described previously (Worliczek et al., 2013), suspended in culture medium and applied to the host cells in a density of sporozoites:host cells of 1:30. The infected cell cultures were incubated at 37°C, 5% CO_2_ and culture medium was exchanged at 1, 4, and 7 days post infection (dpi).

### 2.3 *S. neurona* culture and immunofluorescence assays

*S. neurona* strain SN3 was cultured, transfected and processed for immunofluorescence as described before (Dubey et al., 2017; Howe et al., 2018). In short, parasites were cultured and maintained in bovine turbinate (BT) cell monolayers. For IFA, extracellular merozoites were used to infect BT cells in 24-well plates containing coverslips. Typically, on day 3 post infection, infected BT cell monolayers were methanol-fixed for downstream immunofluorescence experiments. Freshly isolated extracellular *S. neurona* merozoites were used for transient expression of YFP-tagged TgIMC15 (Anderson-White et al., 2011).

### 2.4 *B. bigemina* immunofluorescence assays

*B. bigemina* indirect immunofluorescence assays were performed on >10% parasitemia bovine red blood cells by air drying drops in 10-well slides (Electron Microscopy Sciences), followed by 100% methanol fixation for 5 min at -20°C, 3 rinses in PBS and blocking in 3% BSA in PBS for 1 hr at RT. Primary antibodies diluted in blocking solution (rabbit TgCentrin1 1:4000 (Fung et al., 2012), mouse MAb 12G10 α-tubulin 1:100 (University of Iowa Hybridoma Bank) (Jerka-Dziadosz et al., 1995); mouse ascites MAb B-5-1-2 α-tubulin 1:200 (Sigma-Aldrich #T5168); rat TgEB1 1:100 (Chen and Gubbels, 2019); guinea pig BbIMC1a 1:500) were incubated for 1 hr at RT, followed by 3 washes for 5 min in PBS, Alexa488 or 594 conjugated secondary antisera (ThermoFisher) diluted 1:400 in 0.1% BSA in PBS for 45 min at RT followed by 1 wash for 5 min in PBS containing 1.5 µg/ml 4’,6-diamidino-2-phenylindole (DAPI) to stain DNA, and two additional 5 min washes with PBS before mounting with Fluoro-Gel (Electron Microscopy Sciences). A Zeiss Axiovert 200 M wide-field fluorescence microscope was used to collect images, which were deconvolved and adjusted for phase contrast using Volocity software (Quorum Technologies). SR-SIM was performed on a Zeiss ELYRA S.1 system in the Boston College Imaging Core in consultation with Bret Judson. All images were acquired, analyzed and adjusted using ZEN software and standard settings.

### 2.5 *C. suis* indirect immunofluorescence assays

*C. suis* indirect immunofluorescence assays were performed on infected IPEC-J2 cells grown on glass cover slips as described above at dpi 7 and 8. Infected cells on cover slips were washed in PBS, fixed with ice cold methanol for 10 min and subsequently washed twice with PBS. Blocking was performed for 20 min with Superblock® T20 (PBS) blocking buffer (Thermo Scientific) with 1% (v/v) normal goat serum at RT. Primary antibodies (guinea pig TgNuf2 (Farrell and Gubbels, 2014); mouse mAb 6-11B-1 acetylated α-tubulin, Invitrogen) were diluted to 1:1000 (v/v) in blocking solution, and incubated for 90 min at 37°C, followed by 2×10 min washes with PBS-Tween 20 (0.01%, v/v), incubation with secondary antibodies diluted 1:600 in blocking solution for 40 min at 37°C, 2×10 min washes with PBS-Tween 20, followed by 1 wash for 5 min in PBS containing 1 µM DAPI, 4 washes of 5 min at RT with PBS and quick immersion in ddH_2_O, before mounting with Aqua Polymount (Fisher Scientific). Mounted samples were hardened for 2-5 days before imaging. A Zeiss LSM 510 Meta with a 63x plan apochromat oil immersion objective was used to collect Z-stacks using ZEN 2009 Light Edition (Carl Zeiss Microimaging GmbH, Jena, Germany). Z-stacks were subsequently deconvoluted using Huygens Essential 4.3 software (Scientific Volume Imaging Inc., The Netherlands) with measured point spread functions (PSF) for the respective channels. Z-projections (maximum intensity projections of split channels) were computed with ImageJ 1.48e (National Institutes of Health, Bethesda, MD, USA).

### 2.6 Transmission electron microscopy

For thin-section transmission, pellets of >10% parasitemia *B. bigemina* infected bovine red blood cells were fixed with 2.5% glutaraldehyde in 0.1 M sodium cacodylate (Sigma) buffer (pH 7.4) for 1 hr at RT and processed as described (Nishikawa et al., 2005) before examination with a Philips CM120 Electron Microscope (Eindhoven, The Netherlands) under 80 kV.

Pellets of >10% parasitemia *B. divergens* infected human red blood cells were fixed with 2.5% glutaraldehyde 1.25% paraformaldehyde and 0.03% picric acid in 0.1 M sodium cacodylate buffer (pH 7.4) overnight at 4°C. Cells were washed in 0.1 M cacodylate buffer and post fixed with 1% OsO_4_/1.5% KFeCN_6_ for 1 hr, washed 2x in water, 1x maleate buffer (MB) 1x and incubated in 1% uranyl acetate in MB for 1 hr followed by 2 washes in water and subsequent dehydration in grades of alcohol (10 min each; 50%, 70%, 90%, 2×10 min 100%). The samples were then put in propyleneoxide for 1 hr and infiltrated overnight in a 1:1 mixture of propyleneoxide and TAAB (TAAB Laboratories Equipment Ltd). The following day the samples were embedded in TAAB Epon and polymerized at 60°C for 48 hrs. Ultrathin ∼60 nm sections) were cut on a Reichert Ultracut-S microtome, picked up on to copper grids stained with lead citrate and examined in a JEOL 1200EX Transmission electron microscope and images were recorded with an AMT 2k CCD camera.

### 2.7 Phylogenetic analysis

The following 18S rRNA gene sequences were used: *Cryptosporidium parvum* (GenBank accession no.: L25642.1), *Cryptosporidium muris* (AB089284.1), *Babesia bovis* (L19077.1), *Babesia bigemina* (JQ437261.1), *Babesia divergens* (AJ439713.1), *Babesia duncani* (HQ285838.1), *Babesia microti* (AB219802.1), *Theileria annulata* (EU083801.1), *Theileria parva* (L02366.1), *Theileria orientalis* (HM538200.1), *Cytauxzoon felis* (L19080.1), *Theileria equi* (KM046918.1), *Hepatozoon canis* (KX712124.1), *Plasmodium falciparum* (XR_002966654.1), *Plasmodium berghei* (XR_002688202.1), *Plasmodium vivax* (XR_003001225.1), *Toxoplasma gondii* (L24381.1), *Hammondia hammondi* (KT184369.1), *Neospora caninum* (U03069.1), *Besnoitia besnoiti* (KJ746531.1), *Cystoisospora suis* (KF854251.1), *Sarcocystis neurona* (KT184371.1), *Hyaloklossia lieberkuehni* (AF298623.1), *Goussia janae* (GU479644.1), *Goussia neglecta* (FJ009242.1), *Cyclospora cayetanensis* (XR_003297358.1), *Eimeria callospermophili* (JQ993648.1), *Eimeria tenella* (KT184354.1), *Eimeria necatrix* (KT184349.1), *Eimeria maxima* (KT184346.1), *Eimeria brunetti* (KT184337.1), and *Eimeria acervulina* (KT184333.1). Sequences were aligned using Clustal Omega (Sievers et al., 2011) and a consensus phylogenetic tree was generated using Geneious Prime V2019.1.3 (Invitrogen) using the Jukes-Cantor genetic distance model, neighbor-joining tree build method, bootstrapped 100x using *C. parvum* as outgroup.

### 2.8 Generation of *B. bigemima* IMC1a antiserum

We identified five *B. bigemina* IMC proteins by reciprocal BLASTP searches on EuPathDB (Warrenfeltz et al., 2018) with the 14 *T. gondii* alveolin-domain containing IMC proteins (Anderson-White et al., 2011): PiroplasmaDB.org accession number BBBOND_0204530, BBBOND_0401990, BBBOND_0201220, BBBOND_0208040, BBBOND_0300730). *B. bigemina* BBBOND_0204530, which we named BbIMC1a, was most conserved and the whole ORF of 600 bp was synthesized by TwistBioscience (San Francisco, CA), amplified with primers #4831 pAVA-Gibson-Twist-F gaagctcagacccaggg and #4832 pAVA-Gibson-Twist-R tgcagaacttgttcgtgctg to remove the linkers and cloned by Gibson assembly into *Pme*I/*Nru*I digested plasmid pAVA0421 (Alexandrov et al., 2004), expressed as a 6-His tag fusion in *Escherichia coli* BL21-RIPL, purified by Ni-NTA chromatography under denaturing conditions (Invitrogen), refolded, and used to immunize a guinea pig (Covance, Inc). Serum was affinity purified as described previously (Gubbels et al., 2006) against recombinant His6-BbIMC1a.

## 3 Results

### 3.1 External budding

Schizogony is defined as: schizo = split (or cleft); gony = birth (genesis of a class of thing). It is generally understood as meaning “multi-fission” and is applied to division modes producing more than two daughter cells per division by peripheral (or cortical) budding from the plasma membrane of a polyploid, multi-nucleated mother cell (Fig 2). However, confusion enters with the term “schizont”, which is more widely defined and used to describe any polyploid intermediate cell state regardless of division mode (Fig 4-6): schizo = split (or cleft); -ont = a being (from Greek einai to be) i.e., “to be split”. We forewarn that the term ‘schizont’ is and will be used across division modes for polyploid cells, and thus will not always refer to schizogony per se.

**Figure 3.**
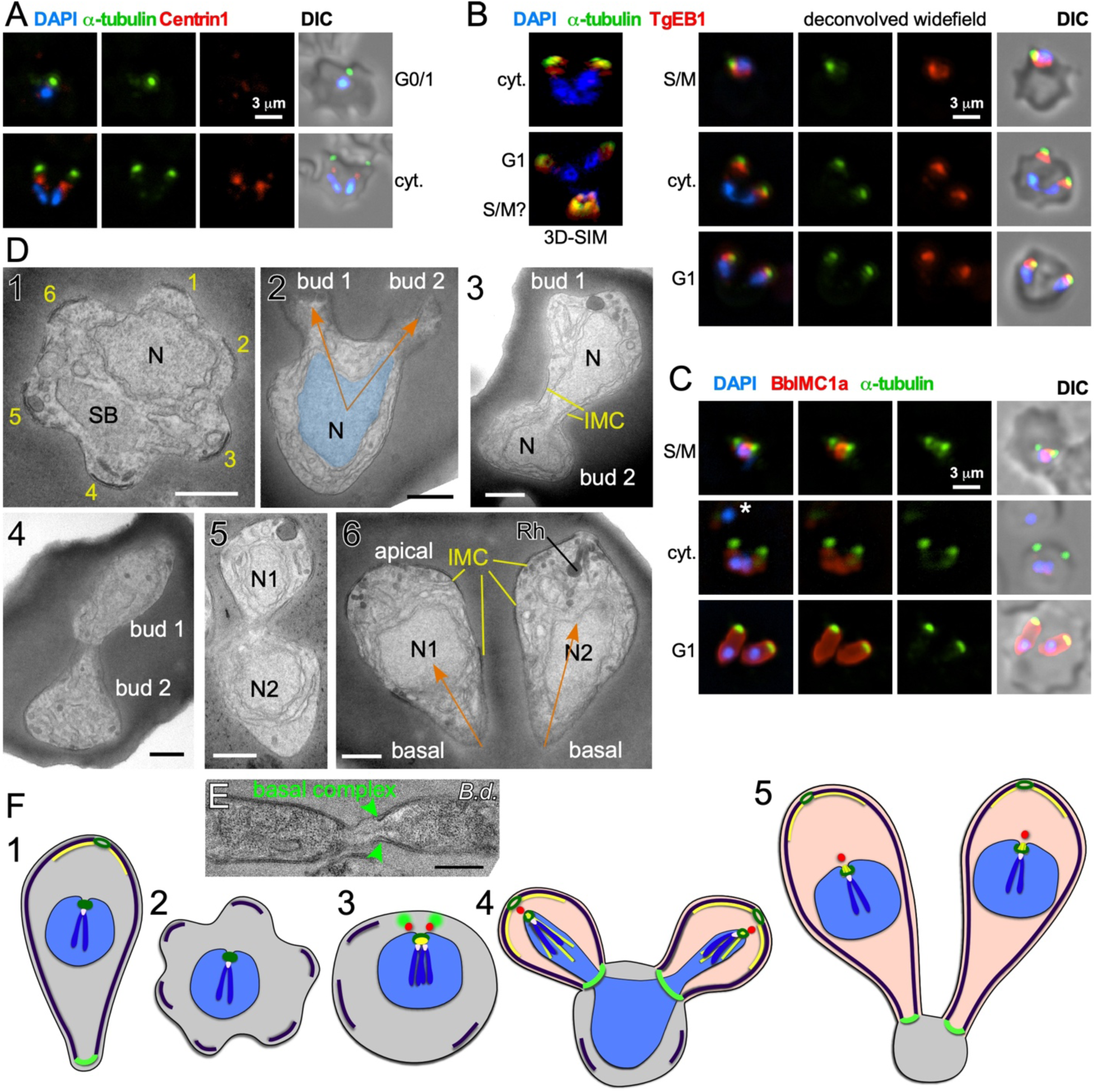
Binary fission by *Babesia* spp. **A-D**. *Babesia bigemina* iRBC stages. **A**. Immunofluorescence using MAb α-tubulin 12G10 (green) and TgCentrin1(red) polyclonal antibody. Centrin staining is not observed in interphase (G1) while diffuse and weak during cell division (cyt.). **B**. Immunofluorescence using MAb α-tubulin 12G10 (green) and microtubule (+)-end binding protein TgEB1 polyclonal antibody (red) demonstrates that the cortical microtubules are relatively short and do not reach the nucleus. **C**. Immunofluorescence using polyclonal guinea pig BbIMC1a antiserum (see Fig S1 for validation) and MAb α-tubulin 12G10 shows the IMC is present during G/M, outlines budding daughters (cyt.), and extends along the length of the mature G1 parasites. * marks a recently invaded parasites wherein the cytoskeleton is completing disassembly; we observed many trophozoites without detectable PbIMC1a. **D**. Transmission electron microscopy of progressive iRBC developmental stages (1→6). Panel 1 is a disassembling merozoite already escaped from the vacuole displaying typical pattern of 6 numbered remnants of the disassembling mother IMC. Panel 2 represents early daughter formation (‘Mickey Mouse’) with the nucleus being separated into the daughter buds. Panels 3, 4, and 5 represent different sections though developing daughters where showing parallel assembly (in contrast to polar assembly) consistent with the angle under which the daughter buds assemble. Panel 6 displays two just divided daughters under the typical angle. N, nucleus, SB spherical body. Yellow marks IMC; orange arrows mark the typical angle and direction of daughter buds. Scale bars represent 500 nm. **E**. *Babesia divergens* basal complex upon completion of cell division. Scale bar represents 500 nm **F**. Schematic of apicomplexan asexual cell division by binary fission, which mechanistically represents a binary form of schizogony. Note that the mother’s cytoskeleton is largely disassembled following successful host cell invasion resulting in a pleomorphic cell (2), but that fragments of the IMC are remaining throughout the division stages (2-4).

**Figure 4.**
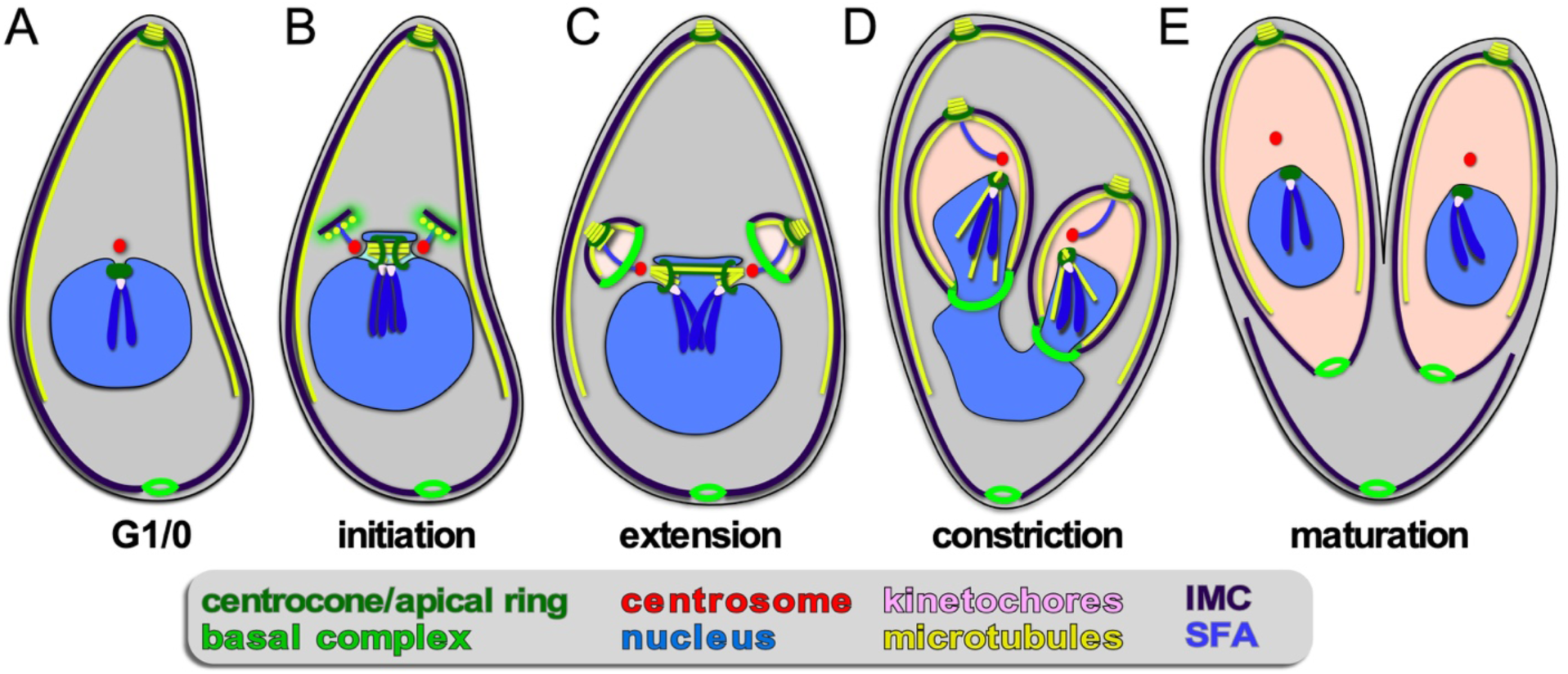
Apicomplexan asexual cell division by endodyogeny. **A-E** show progressive steps of cell division as observed for *T. gondii* tachyzoites. Two daughters bud internally, while the mother’s cytoskeleton is maintained and only destabilized just before emergence of nearly mature daughters (E). A dark blue SFA fiber anchors the centrosomes in the apex of the daughter buds.

#### 3.1.1 Schizogony with karyokinesis

A wide array of Apicomplexa impacting humans or livestock divide by schizogony, such as nearly all Eimeriidae family members of the Coccidian parasites (Dubremetz, 1973; Joyner and Long, 1974; Dubremetz and Elsner, 1979; Ferguson et al., 2008), the genus of *Plasmodium* in the order of Haemosporida (Arnot et al., 2011; Stanway et al., 2011) as well most (but not all!) piroplasms compromising *Theileria* and *Babesia* spp. (Mehlhorn and Shein, 1984) (Fig 1). The piroplasms are tick-transmitted parasites that multiply in red blood cells without forming pigments (order Achromatorida). Both the piroplasms and the *Plasmodium* spp. (order Chromatorida: form pigments in red blood cells) reside in the clade Haemosporidia for their shared residence within red blood cells. Our discussion here is based mostly on *Plasmodium* schizogony in the red blood cells as it is best studied system, however, insights likely apply widely. A defining principle of schizogony is the disassembly of the mother’s cytoskeleton shortly following completion of host cell invasion, resulting in an amoeboid or pleomorphic cell (Hepler et al., 1966; Gruring et al., 2011) (Fig 2A2) which then undergoes several cycles of DNA replication and nuclear division (Fig 2A3,4). These rounds of DNA replication without budding are controlled by cdc2-related kinase CRK4 which resides in the nucleoplasm and is associated with phosphorylation of the DNA replication machinery (Ganter et al., 2017). Interestingly, the mitotic cycles of the nuclei sharing the same cytoplasm are not synchronous. Mechanistically, this observation is anchored in a differential maturation model of mother and daughter centrosomes following their duplication. Although this model has been firmly demonstrated in higher eukaryotes and is defined by proteins in the centrosome’s distal and sub-distal appendages (Bornens and Gonczy, 2014), it has not been formally confirmed in the Apicomplexa. The last cycle of S/M-phase, in contrast, is synchronous and coupled to the synchronous budding of the daughters at the end of schizogony (Ferguson et al., 2008; Kono et al., 2016; Rudlaff et al., 2019) (Fig 2A4,5). Budding follows the activation of the centrosomes’ outer-cores, which is always tied to simultaneous centrosome inner-core activation. The exact timing of the nuclear cycle before the onset of budding has not been clearly resolved. Notably, the level of PfCRK4 drops when the budding cycle is about to start, but it is not clear whether this is the defining signal synchronizing the nuclear cycles and/or activating the budding cycle. Potentially, a diffusible signaling protein could respond to the depletion of nutrients, the accumulation of waste, limited space, or act as a quorum sensor, and subsequently synchronize or halt the nuclear cycle before activating the synchronous budding phase. Since the number of offspring varies across *Plasmodium* species, even between different *P. falciparum* strains, across different development stages (e.g. the liver cell expands and produces up to 90,000 merozoites from infection by a single sporozoite (Vaughan and Kappe, 2017)) there is also a strong, programmed genetic component determining the number of offspring (Reilly et al., 2007). Either way, when the ‘commitment to budding’ checkpoint is cleared, the centrosomes that anchor the spindle pole and chromosomes, reorient to associate with the plasma membrane, a step which is mediated by Cyc1 (Robbins et al., 2017). Note that the centrosomes in *Plasmodium* spp. technically are centriolar plaques as these parasites lack canonical centriolar structures. Following plasma membrane docking of the centrosomes, the daughter cytoskeletons start to assemble in connection with the plasma membrane and the buds move outward (Fig 2A4,5). In *P. falciparum* the maturation and release of daughters is mediated by CINCH, a contractile protein in the basal complex (Rudlaff et al., 2019).

#### 3.1.2 Schizogony with limited karyokinesis

Although not separately recognized in the naming conventions, a variation on the classical schizogony as described in the preceding section occurs during sporozoite formation in the invertebrate vector wherein sexual development takes place (Fig 1). This happens in the mosquito midgut for the *Plasmodium* spp (Simonetti, 1996) and in the tick salivary gland for most piroplasms, including *Theileria* spp. and many *Babesia* spp. (Mehlhorn and Shein, 1984; Jalovecka et al., 2018). *P. berghei* and *P. falciparum* sporogenesis have been most extensively studied, which occurs in an extracellular oocyst residing under the basal lamina of the midgut wall (Simonetti, 1996). Like in classical schizogony, the mother’s cytoskeleton is disassembled resulting in a pleomorphic cell. Here, DNA replication and mitosis are not always followed by karyokinesis leading to a patchwork of nuclei, with varying levels of ploidity (Fig 2B). Again, the individual nuclei are at different levels of S-phase and mitosis, however, within one polyploid nucleus this cycle is largely synchronous (Howells and Davies, 1971b; a; Schrevel et al., 1977; Sinden and Strong, 1978). Typically, a round of sporogenesis produces thousands of individual sporozoites from a single mother cell. Similar to classical schizogony, the budding of daughters from the cortex occurs simultaneously and is coupled to a synchronized round of S-phase and mitosis (Howells and Davies, 1971a; Schrevel et al., 1977; Araki et al., 2019; Pandey et al., 2019). This process is conserved in *Theileria* and *Babesia* spp. as well, where multiple and multi-lobed nuclei are especially prominent (Moltmann et al., 1983). However, some *Babesia* spp., produce sporozoites through binary fission (Mehlhorn and Shein, 1984).

In summary, the intriguing phenomenon is that karyokinesis seems to be optional in these polyploid schizonts. The synchronized cycles of mitosis within a single nucleus suggest that mitotic cycles are organized on the nucleoplasm level just as in classical schizogony, whereas commitment to budding is a factor shared across the whole cytoplasm. How these events are controlled at the molecular level has not been determined.

#### 3.1.3 Binary and multiple fission

Binary fission is defined as “the formation of two daughter cells per division round”. Among the Apicomplexa, binary fission is used to describe the division process in the red blood cell stages of many *Babesia* and *Theileria* parasites. Most *Babesia* spp. form two daughters per division round, i.e. classical binary fission. However, in the red blood cell cycle several clades of *Babesia*, including *Babesia duncani* (Conrad et al., 2006) and *B. microti* (Rudzinska, 1981), form four merozoites per division round (Maltese cross), as do all *Theileria* spp. (Conrad et al., 1985; Conrad et al., 1986; Fawcett et al., 1987; Uilenberg, 2006). This process is known as both schizogony and ‘multiple fission’, however it is generally considered distinct from schizogony, which as mentioned above, is a term historically used when many more daughters are produced. The term “binary fission” does not imply that budding is involved and does not intuitively connect with “multiple fission”, which produces only four daughters per division round, as defined in schizogony. Due to the lack of clarity on how these processes are related and defined cell biologically, the exact nature of *Babesia* cell division in the red blood cell remains to be elucidated. The ambiguity of the used terms (i.e. binary fission, budding, schizogony) to describe this process further exacerbates this confusion. In a 1978 publication study using transmission electron microscopy (TEM) the interpretation was as follows (Potgieter and Els, 1977): “The trophozoites were surrounded by a single membrane, were pleomorphic in shape and contained large inclusions of host cell cytoplasm, but no cytostomes or food vacuoles could be identified. Reproduction took place through a process resembling schizogony resulting in the production of two merozoites, the cytoplasmic constituents of the original trophozoite (mother cell) being virtually entirely incorporated into the daughter cells in the process”. In essence, the pleomorphic/ameboid trophozoite in combination with the schizogony reference suggest that the process is conceptually very similar to schizogony, except that term does not capture it since only two daughters are formed. Here we combine existing data with new insights from *in vitro* cultivated *Babesia bigemina* to clarify the mechanistic murkiness surrounding binary fission.

*B. bigemina* is a cattle-infecting, tick-transmitted apicomplexan causing vast economic losses (Bock et al., 2004; Suarez et al., 2019). From an experimental point of view, *B. bigemina* cell division is uncomplicated in that it only produces two daughters per division round, and it is a relatively large *Babesia* spp., making it an ideal candidate for microscopy. We focused on the key organizers of cell division: the centrosome (or centriolar plaques) and cortical cytoskeleton. *T. gondii* Centrin1 antiserum on *B. bigemina* showed a specific, albeit relatively weak, signal, but only during daughter budding (Fig 3A). It therefore appears that centrosome composition is dynamic, which was recently also reported for *P. berghei* Centrin4 as this signal was only on the centrosome during mitosis and budding, but not in mature or recently invaded parasites (Roques et al., 2019). In contrast, *Plasmodium* Centrin3 is a popular marker of centrosomes and is always associated with the centrosome, indicating that the dynamics of different Centrins can vary (Mahajan et al., 2008; Kono et al., 2012). We detected microtubules with α-tubulin MAb 12G10 generated against ciliate *Tetrahymena thermophila* (Jerka-Dziadosz and Frankel, 1995), which has shown broad reactivity in the Apicomplexa. A spot focused toward the apical end was observed (Fig 3A), but we never observed tubulin in the nucleus, suggesting this antibody does not detect the mitotic spindle. In *Plasmodium* the spindle was successfully visualized with monoclonal antibody B-5-2-1 generated against sea urchin α-tubulin (Gerald et al., 2011), but in *bigemina* we observed the same patterns as for MAb 12G10 (not shown). In *T. gondii*, detecting the spindle microtubules has also been challenging, but a polyclonal serum against TgEB1, a microtubule (+)-end binding protein, was very specific for the spindle (Chen et al., 2015b). Reactivity of α-TgEB1 in *B. bigemina* across all parasite stages highlighted a spot basal of the apical tubulin staining (Fig 3B). We interpret that EB1 does not bind to the spindle microtubules in *B. bigemina*, but instead associates with the (+)-end of the subpellicular microtubules emanating from the MTOC at the very apical end. This also indicates these microtubules are relatively short and only cover the very apical cap of the parasite. To better visualize the cortical cytoskeletal scaffolds that drive budding we tested polyclonal antisera generated against *T. gondii* IMC proteins (Gubbels et al., 2004; Anderson-White et al., 2011), but unfortunately none showed a signal. Therefore, we generated a specific *B. bigemina* IMC antiserum against BBBOND_0204530, the closest relative to the *T. gondii* alveolin domain containing IMC proteins (Anderson-White et al., 2011), which we named BbIMC1a (Fig S1). IFAs with α-PbIMC1a nicely highlighted the cortical cytoskeleton during budding, extending basally beyond the microtubule signal. In mature parasites we observed signal along the entire length of the merozoite (Fig 3C). In trophozoites we also observed parasites identified by their DAPI signal with weak or variable intensities of either the IMC1a or tubulin signals that appeared disorganized, or at least not to outline a merozoite (Fig 3C top panel and middle panel marked by an asterisk). We interpreted these amorphic signals to represent trophozoites that are at the stage between invasion and the start of cell division. However, across all parasites in division we observed parasites forming next to each other, with apical ends emerging first with V-shape symmetry.

To obtain a higher resolution of the mechanistic steps in the *B. bigemina* division process we performed TEM. We focused on stages after the parasites escaped from their vacuole and entered G1 growth phase (trophozoites). We observed a high frequency of zoites wherein the IMC breaks into 5-6 pieces, still in curved structures on the edge consistent with an ameboid or pleomorphic status for the zoites (Fig 3D1). This consistent observation suggests that disassembly of the mother cytoskeleton is spatially organized resulting in a symmetrical appearance, which reflect the trophozoites identified by IFA (Fig 3C). Similar observation on disassembling mother IMC have been reported for *Plasmodium* ookinetes in the mosquito midgut (Carter et al., 2007), and for sporozoites in liver cells, where regularly spaced breaks occur in the IMC, although not quite as symmetrically organized as seen here in *B. bigemina* (Jayabalasingham et al., 2010). In a budding zoite two daughter buds are emerging on one side of the cell (like rabbit ears), and not on polar opposites (Fig 3D2). The shape of the nucleus is also consistent with the anchoring of the nucleus at the apical end of the daughters, which, by using other division modes as a guide, we assume is through the spindle poles (Potgieter and Els, 1977). For the growing stages, we captured many transverse cross sections of two side-by-side budding daughters with the nucleus at various stages of division (Fig 3D 3-5). Finally, completely formed daughters appear side by side under a similar angle as the early daughter buds (compare the orange arrows in Fig 3D2 with 3D6). Additional TEM images of *Babesia* buds in a V orientation consistent with our data have been reported elsewhere (Friedhoff and Scholtyseck, 1977; Potgieter and Els, 1977; Scholtyseck, 1979). A section through the basal ends of still connected *Babesia divergens* parasites (predominantly a cattle parasite but opportunistic in humans) displays an electron dense structure on the basal end of the IMC consistent with a basal complex (Fig 3E). Although we did not capture a section through this structure in *B. bigemina*, a similar appearance of the basal complex in *B. bigemina* has been reported previously (Potgieter and Els, 1977).

Overall, the budding of daughters in *B. bigemina* shares many features seen across asexual apicomplexan development (Fig 3F). First, the angle between the daughter buds fits with the pleuromitosis (closed mitosis with the spindle poles in close proximity at one side of the nucleus) model dictated by closely apposed centrosomes at the apically defining side of the nucleus. This is observed across the Apicomplexa that have been studied (Gerald et al., 2011; Francia and Striepen, 2014). As a result, daughter buds are formed next to each other rather than in opposing orientations (middle panel Fig 3F3-5). The *B. bigemina* trophozoite has an ameboid appearance after invasion and cytoskeleton is disassembled (Fig 3E1, 3F2; and (Potgieter and Els, 1977)), which likewise is observed during the corresponding stage of *P. falciparum* (Gruring et al., 2011). Taken together, these data firmly illustrate that the processes known as binary fission and schizogony, and by extension multiple fission as seen in *Theileria* spp., are mechanistically the same and only differ in the number of nuclear replication cycles before the onset of cytokinesis. Collectively, we now group these division forms together under the umbrella “external (aka cortical) budding”

#### 3.1.3 Other variations on cortical budding

Schizogony by cortical budding as described in the preceding sections are found in various asexual stages. Other forms on cortical budding can be found in sexual development stages, which we will briefly highlight here to illustrate the spectrum of division modes. *Plasmodium* male gametocytogenesis produces flagellated microgametes in a process known as “exflagellation”. This process occurs in the red blood cell and unfolds by several fast rounds of S/M-phase resulting in a single polyploid nucleus (Sinden et al., 1978; Sinden, 1983). The cortical ‘budding’ of microgametes, which at the cytoskeletal level basically are a single flagellum and only contain microtubules, progresses while the nuclear material partitions. Interestingly, a recent study using Ndc80 as a marker for the kinetochores demonstrated that the genome size universally increases to 8N, but that chromosome replication is asynchronous (Pandey et al., 2019). This is in contrast to what is observed during asexual schizogony, where the rule is that the mitotic cycles are synchronous within a shared nucleoplasm.

In addition, microgametocyte formation of *T. gondii* also progresses through a cortical budding process (Ferguson et al., 1974). As discussed in detail below, it is salient to note that the asexual division of *T. gondii* is by internal budding, not through cortical budding. *T. gondii* microgametocytes formation plays out by the association of 1N nuclei in a multi-nucleated microgamont with plasma membrane of the mother cell. Interestingly, the mother cell (macrogamont) maintains an IMC where in holes are present to facilitate association of the nuclei with the plasma membrane. From here, bi-flagellated microgametes bud outward in a cortical budding process (Ferguson et al., 2008). Interestingly, this process is conserved across the Coccidia, as it is also reported for *Eimeria* spp. which, in contrast to *T. gondii*, replicate asexually by schizogony (Ferguson et al., 1980) (Fig 1). Taken together, the notable features here are maintenance of the cortical cytoskeleton (Dubey et al., 2017) in combination with cortical budding while S-phase, mitosis and karyokinesis have been completed before the onset ‘budding’.

### 3.2 Internal budding

#### 3.2.1 Endodyogeny

Endodyogeny is defined as: endo = inner/internal; dyo = two; geny = genesis, meaning birth; production/generation/origin. This term is applied to a type of reproduction in which two daughters are formed within a parent cell (Goldman et al., 1958; Sheffield and Melton, 1968). Endodyogeny has been described in detail for *T. gondii* tachyzoites (Nishi et al., 2008; Anderson-White et al., 2012) and is employed as well by its closest evolutionary neighbors comprising several genera of the Sarcocystidae (Fig 1, 4). These include *Hammondia, Neospora* and *Besnotia* spp. as well as the tissue cyst-forming stages of *Sarcocystis neurona*. During endodyogeny, the mother’s cytoskeleton is not disassembled following completion of invasion, and two daughter buds assemble on centrosomes residing within the cytoplasm. Only at the very last stage of daughter budding is the mother’s cytoskeleton disassembled and is the plasma membrane deposited on the new daughters, which is mediated by recruitment of the “gliding motor complex” to the IMC. This complex contains a multi-acylated protein glideosome associated protein, GAP45, that is anchored in both the plasma membrane and the IMC outer membrane and ‘zippers’ these structures together in an apical to basal direction (Gaskins et al., 2004; Frenal et al., 2014). Many of the cell cycle checkpoints throughout endodyogeny have been resolved. Specifically, two checkpoints have been described upon commitment to mitosis and budding, one likely dedicated to mitosis and the other to budding. This is thought to facilitate the uncoupling of S/M cycles from budding in the multi-daughter division modes and differentially activate the centrosome inner- and outer-cores (Suvorova et al., 2015; Naumov et al., 2017; White and Suvorova, 2018). Like in schizogony, daughter budding occurs in sync with S/M-phase and karyokinesis. Upon emergence of daughters, a narrow cytoplasmic bridge at the basal end remains connected with a residual body containing remnants of the mother cell, which will be largely resorbed into the daughters (Frenal et al., 2017b; Periz et al., 2019). As a result, the daughter parasites remain in contact with each other and could therefore have the false appearance of endopolygeny. To differentiate these two processes, this division process has been referred to as ‘repeated endodyogeny’.

#### 3.2.3 Endopolygeny

Endopolygeny is defined as: endo = inner/internal; poly = multiple; geny= genesis/birth. This term is used to describe the reproduction types wherein more than two individuals are formed simultaneously within the cytoplasm (i.e. not from the cortical periphery) of a polyploid parent cell. One of the key features of internal budding is that the mother’s cytoskeleton is maintained throughout cell division and only dissembles just before the emergence of almost completely assembled daughters. Although not differentiated in the naming conventions, two sub-forms can be distinguished: either the polyploid mother cell can be multinucleate, or it can contain one large polyploid nucleus depending on whether karyokinesis follows each round of S-phase. Both these forms of budding are found within the tissue cyst-forming Coccidia, more specifically the Sarcocystidae (Fig 1, 5, 6). These division forms are related to endodyogeny, since the mother’s cytoskeleton is maintained throughout the division cycle until the maturation of the daughter cells. We define these three forms here collectively as “internal budding”. Internal budding by the Sarcocystidae is unique among the Coccidia as other families, notably the closely related Eimeriidae, replicate by schizogony (Fig 1). Two emerging systems exist to study the variations in progression of endopolygeny: *Sarcocystis neurona* is a model for studies representing the form without completing karyokinesis (Fig 5) (Vaishnava et al., 2005; Dubey et al., 2017), whereas merogony of *Cystoisospora suis* provides an accessible *in vitro* model for the form including karyokinesis after each S/M-round (Fig 6) (Worliczek et al., 2013).

**Figure 5.**
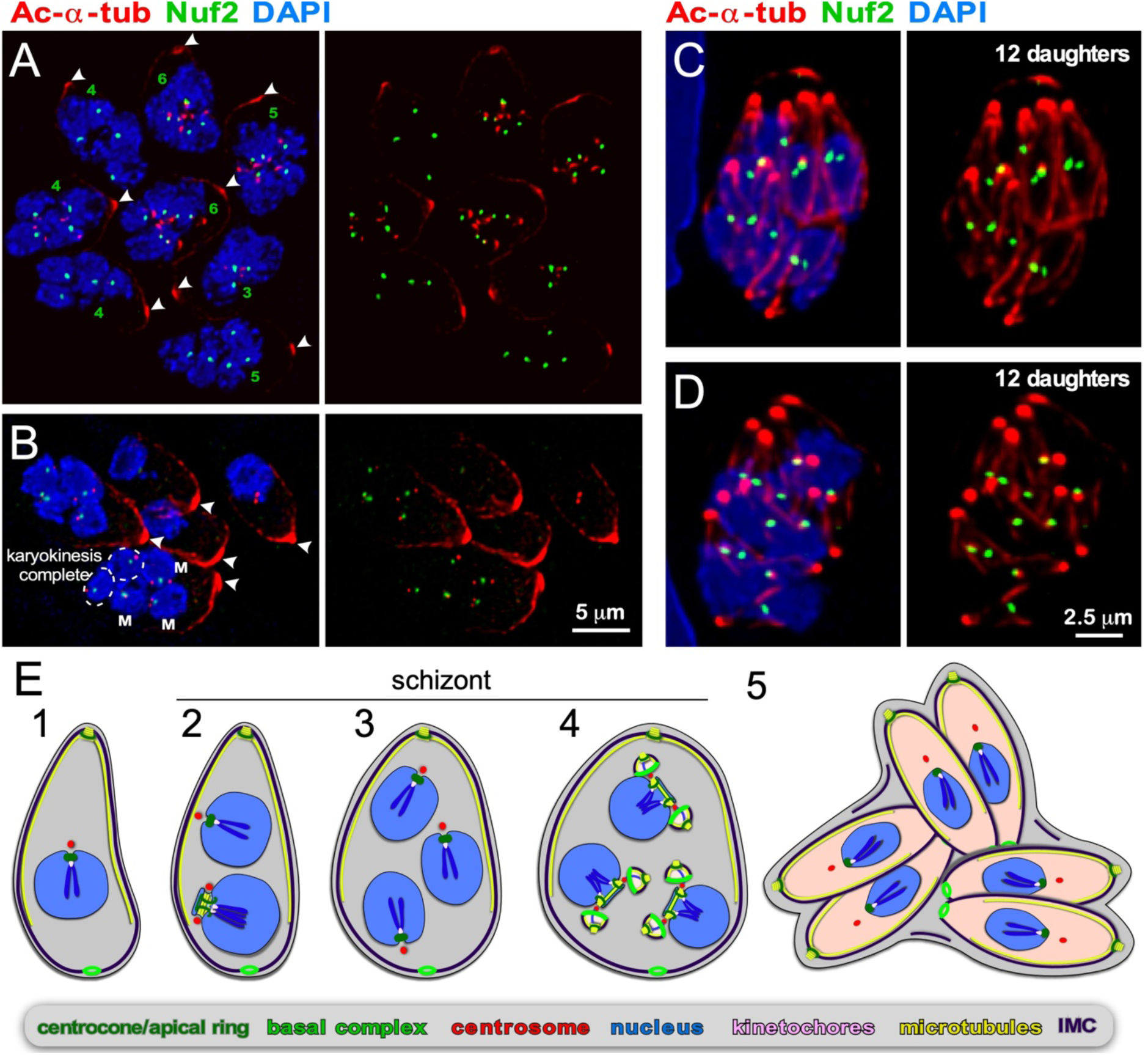
Endopolygeny with karyokinesis during merogony of *Cystoisospora suis*. **A-D**. Parasites at different stages of endopolygeny. Acetylated (Ac) α-tubulin (red) marks the mother cells conoid (arrow heads in A, B), subpellicular microtubules, the spindle poles associated with nuclei undergoing mitosis, and the daughter conoids and subpellicular microtubules (C, D). Antiserum generated against TgNuf2 (green) marks the clustered kinetochores at mitotic spindles. A. Eight parasites at different stages in the division cycle. Green numbers indicate the number of spindle poles seen per mother cell, which is expanding non-geometrically in five of the parasites indicating asynchronous nuclear cycles. **B**. Five parasites at different stages of endopolygeny are shown. The parasite on the bottom has five nuclei: nuclei marked “M” are in early stages of mitosis as 2 spindle poles flanking a single 2N kinetochore cluster are seen; the circled nuclei represent just completed mitosis as the kinetochore clusters are relatively weak intensity (1N vs. 2N) and associated with only a single spindle pole. **C, D**. Two examples of cells at the synchronous daughter budding stage, which both produce 12 daughters as enumbered by the number of kinetochore clusters, again, consistent with non-geometric expansion of the nuclei. **E**. Schematic of apicomplexan asexual cell division by endopolygeny with karyokinesis. Note that the mitotic cycles of the nuclei in the same cytoplasm are not synchronous (2) resulting in non-geometrically expanded daughter numbers (4-5). Also note that the mother’s cytoskeleton is maintained throughout the meront (technically ‘schizont’) stages and is only destabilized just before emergence of nearly mature daughters (5).

**Figure 6.**
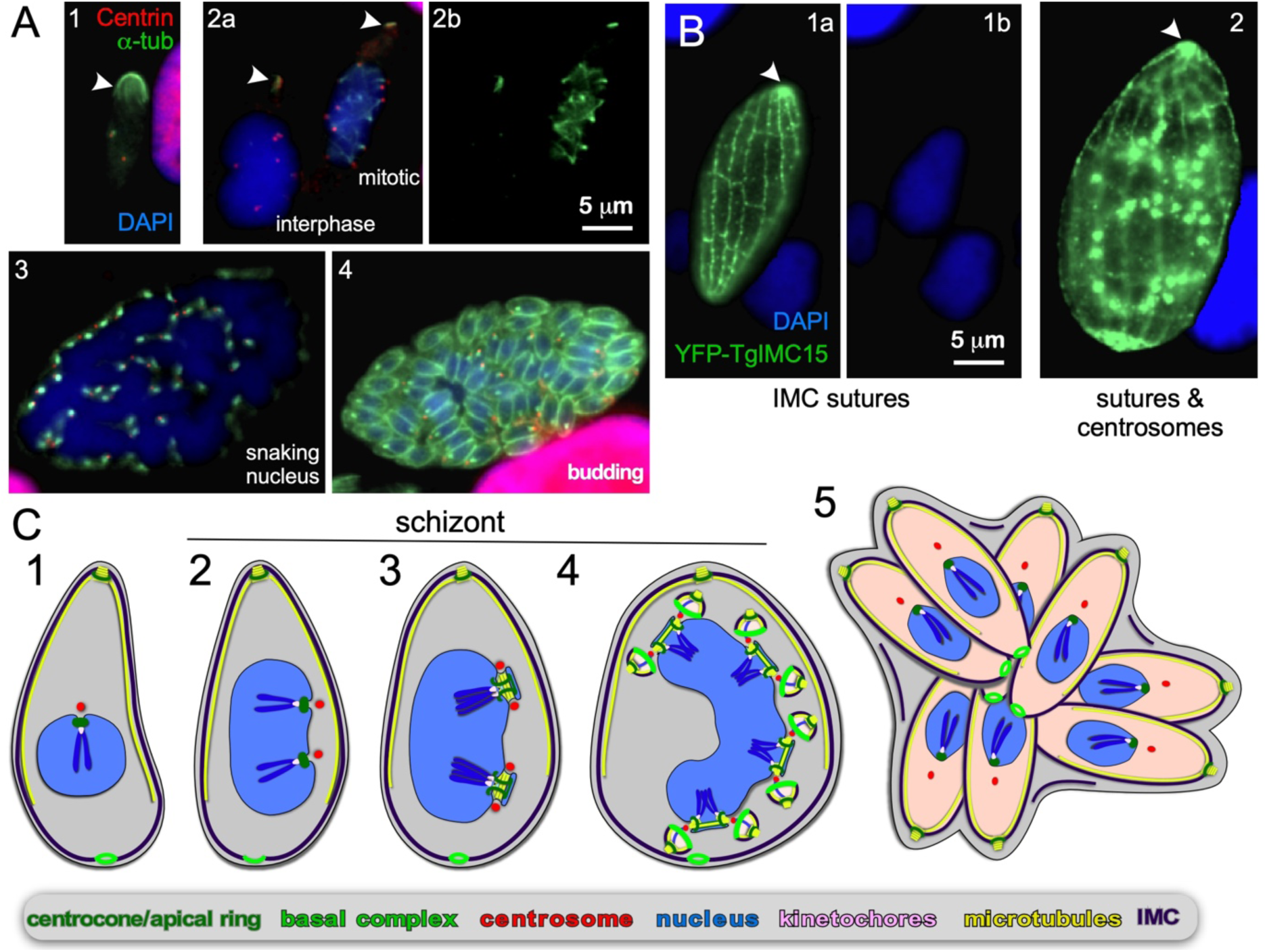
Endopolygeny without nuclear fission by *Sarcocystis neurona* in the intermediate host. **A**. Staining of progressive cell division cycle stages (1-4) with α-tubulin (green) marking the subpellicular microtubules of the mother cytoskeleton (arrow heads), the mitotic spindles (2b, right parasite; 3) and daughter merozoites subpellicular microtubules (4). Centrin staining (red) marks the centrosomes, which due to z-stack selection are not visible for all spindles/parasites. Scale bar applies across panels. **B**. Overexpression of a *Toxoplasma* YFP-IMC15 fusion protein (green) highlights the mother cell’s cortical IMC sutures in both panels, whereas in further progressed panel 2 the bright internal spots mark the centrosomes poised for budding. Arrowhead marks the apical end of the mother parasite. Panel B modified from (Dubey et al., 2017). Scale bar applies across panels. **C**. Schematic of apicomplexan asexual cell division by endopolygeny without karyokinesis. Note the polyploid nuclei undergoing synchronized cycles of M-phase and mitosis resulting in geometrically expanded daughter cell numbers. Also note that the mother’s cytoskeleton is maintained throughout the schizont stages (2-4) and is only destabilized just before emergence of nearly mature daughters (5).

##### 3.2.3.1 Endopolygeny with karyokinesis

*suis* is the causative agent of suckling piglet coccidiosis (Shrestha et al., 2015). Although phylogenetically *C. suis* falls within the cyst-forming Coccidia (Fig 1), tissue cysts have never been observed and the known coccidian development cycle is monoxenic in pigs: the absence of tissue cyst forming capacity is likely a secondary loss in this species (Stuart et al., 1982; Shrestha et al., 2015). *C. suis* asexually replicates in epithelial cells of the porcine small intestine by endodyogeny and endopolygeny, depending on the generation of asexual stages (alternatively called types: meronts/merozoites type I, II and subtype II (Matuschka and Heydorn, 1980). Early asexual division after infection of the gut epithelium is restricted to endodyogeny (often in consecutive cycles within one parasitophorous vacuole, i.e. ‘repeated endodyogeny’), whereas endopolygeny is described from day 3-4 post infection onwards *in vivo* (Lindsay et al., 1980; Matuschka and Heydorn, 1980) and from day 7 onward *in vitro* (Worliczek et al., 2013), concurrent with cells replicating by endodyogeny.

As for *B. bigemina*, we tracked *in vitro* progression of *C. suis* development using the toolbox of reagents we established for *T. gondii*. In *C. suis* zoites undergoing endopolygeny, the mother’s subpellicular microtubule cytoskeleton is clearly visible in large polyploid cells as an apically concentrated microtubular accumulation joined in the mother’s conoid (Fig 5A-D). This confirms the mother’s cytoskeleton is maintained throughout endopolygeny. To track the progression of S/M-phase across the nuclei we used the kinetochore component Nuf2 (Farrell and Gubbels, 2014) in combination with acetylated *α*-tubulin, which marks the spindle poles during mitosis (note that spindle microtubules disassemble during interphase (Farrell and Gubbels, 2014; Chen et al., 2015b)). The first observation is that the status of mitosis (spindle visible by tubulin stain) and the status of kinetochore separation varies among nuclei within the same cell (Fig 5A, B). Furthermore, it is evident that parasites expand non-geometrically, as counted by spindle poles per nucleus varying from 3, 4, 5, to 6 are seen in Fig 5A and B, whereas 12 budding daughters, i.e. non-geometrically expanded, can be discerned in Fig 5C, D. These two observations indicate that the nuclear division cycles are not synchronous, which is consistent with expansion numbers described in the literature (e.g. (Lindsay et al., 1980; Matuschka and Heydorn, 1980)). Here we show for the first time that this is associated with asynchronous nuclear replication cycles. Finally, we show that the final round of mitosis is synchronous for all nuclei and is coupled with daughter budding, as seen in schizogony. Our insights are summarized in the schematic of Fig 5E. Based on the details revealed here, we conclude that like in schizogony, the S/M-phase progression are controlled at the nuclear level, most likely differing maturity of the mother and daughter centrosomes, but that the commitment to budding is synchronized across the cytoplasm.

Reports on *T. gondii* endopolygeny suggest geometric expansion of nuclei: 8–16 progeny per endopolygeny replication cycle in the cat gut have been observed (Ferguson et al., 1974). Historically, the sexual cycle of *T. gondii* has been poorly experimentally accessible, but *in vitro* completion of the sexual cycle in cat intestinal organoids was recently reported, which is expected to provide experimental accessibility (Martorelli Di Genova et al., 2019). To date, *T. gondii* endopolygeny studies have not been so comprehensive to support a strong conclusion in this matter.

An additional notable observation in Fig 5A is the presence of multiple parasites within a single vacuole undergoing endopolygeny. It has been described before that several multinucleate meronts can be present in a single vacuole, which indicates that more than one asexual division occurs in the same cell (Lindsay et al., 1980). As such, this reveals that multiple rounds of budding within the same vacuole are not exclusively found during endodyogeny as described above for *T. gondii*. This phenomenon therefore highlights that the timing of budding is determined by a genetic program, and not controlled by environmental factors, which we also concluded for schizogony.

##### 3.2.3.1 Endopolygeny without karyokinesis

*S. neurona* has a sylvatic cycle in the Americas with small mammals as intermediate hosts and the opossum as the definitive host. However, accidental infection of horses can cause equine protozoal myeloencephalitis (EPM) (Reed et al., 2016). Using the same strategy as mentioned above, we observe an apical concentration of microtubules across cells with progressively larger single nuclei indicating the mother’s cytoskeleton is maintained throughout endopolygeny (arrowheads in Fig 6A). Furthermore, the *S. neurona* IMC can be visualized through overexpression of TgIMC15-YFP, which highlights the sutures in the IMC of the mother throughout development (Fig 6B) (Dubey et al., 2016). When the cell prepares for budding IMC15 also appears on the centrosomes at which point the mother’s IMC is still prominently present (Fig 6B2). Only at the conclusion of the budding process is the mother’s cytoskeleton disassembled, which mimics the dynamics of the IMC during *T. gondii* endodyogeny (Anderson-White et al., 2011; Dubey et al., 2016).

In contrast, to *C. suis*, the S/M cycles of *S. neurona* are completely synchronized: in Fig 6A2 we observe a cell in interphase characterized by absence of spindle microtubules, whereas the right cell displays spindle microtubules associated with each centrosome. This confirms previous observations (Vaishnava et al., 2005), and indicates that when the nucleoplasm is shared, the individual status of centrosome maturation is overruled, likely by a factor diffusing in the nucleoplasm. Late in development the large polyploid 32N nucleus shifts into a multi-lobed, ‘serpentine’ morphology (Fig 6A3), which sets the stage for a final round of S/M-phase now coupled to karyokinesis and internal budding resulting in 64 daughters (Fig 6A4). Thus, when the S/M-phase are not followed by karyokinesis the nuclear cycles remain synchronized and progeny number are the result of geometric expansion.

#### 3.2.4 Budding without multiplication and multiplication without budding

The spectrum of division modes not yet covered spans two more manifestations in sexual development stages. The first, budding without multiplication, is the formation of kinetes in the Heamosporidia. The *Plasmodium* spp. generate ookinetes from a fertilized zygote (ploidity of 2-4N in a single nucleus) that cross the mosquito gut wall (reviewed in (Angrisano et al., 2012; Bennink et al., 2016)). Although the zygote lacks a cortical cytoskeleton, the ookinete has a complete cortical cytoskeleton comprising IMC and cortical microtubules. This cortical cytoskeleton is formed by the ookinete through from the cortex of the zygote in absence of karyokinesis, though a budding stage dubbed ‘retort’ (Canning and Sinden, 1973; Carter et al., 2007). Yet even more exotic variations occur in the piroplasms; both *Theileria* and *Babesia* spp. produce kinetes, which do not bud from the cortex, but into an internal vacuole formed inside the zygote (see (Mehlhorn and Shein, 1984) for a detailed review). Thus, cortical budding can be uncoupled from the nuclear cycle in the zygote stages.

The second variation is multiplication without budding, of which there are three different, possibly related examples. The first is found in the *Babesia* spp. kinetes, which following their formation in the tick gut, cross the midgut and migrate to the tick ovary. Here, the cortical cytoskeleton disassembles and the cell transforms into a pleomorphic cell (Moltmann et al., 1982; Mehlhorn and Shein, 1984). Subsequently, several rounds of S/M-phase without karyokinesis occur to produce a multiploid, lobed nucleus. In a process not understood at neither the mechanistic nor molecular level, these large cells divide into multiple cells, each with a single nucleus. Surprisingly, neither the mother nor the daughter cells have a cortical cytoskeleton. This indicates that at this life stage cell division is independent of any form of budding, and thus indicates that in Apicomplexa the cortical cytoskeleton is not a strict requirement for cell division. From each of these pleomorphic cells a new kinete then forms by budding mediated by cortical cytoskeleton formation into an internal vacuole as described above. The released kinetes then migrate to the salivary glands where they undergo sporogenesis.

The second example of multiplication without budding is a binary variation on the above process found in some *Babesia* spp. (Mehlhorn and Shein, 1984). Kinetes that invaded the tick salivary gland disassemble their cytoskeleton and undergo one round of S/M-phase, karyokinesis and cell division. However, no cortical cytoskeleton is assembled during these binary division rounds. The cortical cytoskeleton is reportedly assembled slowly during the last couple of division rounds resulting in mature sporozoites. Thus, the salient details are that this process is binary and appears to be efficient with resources as the cytoskeleton is only assembled in the last round (unlike endodyogeny, where a complete parasite is assembled each multiplication round). This process has been described for *B. canis* (Schein et al., 1979), a dog parasite, and *B. bovis* (Potgieter and Els, 1976), a cattle parasite, but likely occurs widely across *Babesia* spp. exclusively replicating by binary fission in the red blood cell (Fig 1) (Mehlhorn and Shein, 1984). Thus, across all life stages, these particular *Babesia* parasites seem to have lost the ability to produce more than two daughters per division round, providing a model system to unravel the specifics of the genetic program and regulatory network.

The third example is the process of sporoblast formation in the Coccidia. This occurs within the oocysts released into the environment. Depending on the species, the zygotes can divide themselves into 2 to 4 sporoblasts (this number is a defining feature in diagnosis of fecal oocysts; not dividing is an option as well (Gardiner et al., 1998)). Inside the sporoblasts 2-4 sporozoites form through a budding mechanism coupled to DNA replication. Although studied in many parasites, *T. gondii* produces two sporoblasts and four sporozoites per sporoblast and as such is the simplest form. Consistent with EM data (Ferguson et al., 1979a), it was recently shown that neither microtubules nor IMC proteins were involved in *T. gondii* sporoblast formation (Dubey et al., 2016). Considering the process and life stage, this process therefore appears akin to kinete multiplication in *Babesia* spp., as described above. This thus seems to connect these two parasites across a large phylogenetic distance, which begs the question whether this is an ancestral connection, or a case of convergent evolution.

## 4. Discussion

The picture of cell division across all life stages of the Apicomplexa is that principle differences exist between the sexual and asexual cell division strategies. In the sexual stages, cell number expansion is not necessarily related to budding a daughter cytoskeleton. This shows that budding is in principle not required for apicomplexan cell division, but there are very few mechanistical or molecular details available for these division modes. The other insight is that all invasive zoites with an apical complex form by a budding strategy, which is initiated and coordinated by the centrosome’s outer-core and proceeds in an apical to basal assembly direction. In all the asexual life cycle stages zoite budding is coupled to a complete nuclear cycle (S/M-phase plus karyokinesis). Another general rule across asexual development is that budding is synchronized across the whole cell. This indicates that the centrosome outer-core activation is always coupled to inner-core activation in asexual stages on a shared cytoplasm-wide level (e.g. a diffusible kinase or kinase substrate). The exceptions to this rule are only found in sexual stages: microgametocytogenesis and (oo)kinete formation in the Haemosproridia.

Besides these general rules, there are several variations within asexual division modes which partition into two mechanistically different strategies: internal budding and external budding (Fig 1). Internal budding comprises endodyogeny and the two variations on endopolygeny, whereas external budding captures the two variations on schizogony, binary fission, and multiple fission. The number of offspring in each strategy can vary from two (endodyogeny and binary fission) to several orders of magnitude higher (>10,000 in schizogony). At the furthest extreme are several bovine-infecting *Theileria* spp. of which the schizonts in the white bloods that trigger transformation of their lymphocyte host cells (i.e. leukemia) resulting in division and expansion of the parasites schizont stage along with their host cell (Luder et al., 2009; Chakraborty et al., 2017). As remarked throughout, the number of daughter cells per division round in each life stage is largely genetically controlled, but the details on the controls are just starting to emerge.

Overlapping with the genetic switch committing to budding is the bi-partite centrosome cycle. The apicomplexan centrosome and mitotic cycles are controlled by Nek and Aurora kinases (Reininger et al., 2011; Carvalho et al., 2013; Chen and Gubbels, 2013; Berry et al., 2016; Berry et al., 2018). In *T. gondii* it has been demonstrated that the switch from solely a nuclear cycle to a combined nuclear and budding cycle is controlled by a MAP kinase-like protein (Brown et al., 2014; Sugi et al., 2015; Suvorova et al., 2015). Ultimately, cell cycle progression and the activation of each core is regulated by cyclin and CDK pairs adapted to each apicomplexan cycle, as they likely act independently on the inner- and outer-centrosome cores (Le Roch et al., 2000; Merckx et al., 2003; Alvarez and Suvorova, 2017; Ganter et al., 2017; Naumov et al., 2017; Robbins et al., 2017; White and Suvorova, 2018). An open question is the identity on the factor(s) controlling the pause of nuclear cycles across parasites prior to the final round of coupled S/M-phase and budding in the polynucleate division modes.

The control of the nuclear cycle appears to depend on the ploidy of the nucleoplasm. When each S/M-phase is followed by karyokinesis, then the cycles of individual nuclei are diverging. Mechanistically, this observation is anchored in differential maturation of mother and daughter centrosomes following their duplication; the mother centrosome is sooner primed for another round of replication. This mechanism has been firmly demonstrated in higher eukaryotes and is defined by proteins in the centrosome’s distal and sub-distal appendages (Bornens and Gonczy, 2014). However, this has not been directly demonstrated yet for the apicomplexan centrosome (Morlon-Guyot et al., 2017; Courjol and Gissot, 2018; Chen and Gubbels, 2019). At first sight, contrasting insight comes from synchronized endodyogeny observed in *T. gondii* tachyzoites. Upon completion of cell division, *T. gondii* daughters stay connected through a cytoplasmic actin bridge maintained by Myosin I (Frenal et al., 2017b; Periz et al., 2017). If this bridge is intact, all conjoined tachyzoites undergo endodyogeny in synchrony resulting in geometric expansion numbers. However, the cell division cycles of parasites sharing the same vacuole become uncoordinated if the bridge is disrupted. Strengthening this conclusion is the observation that 4-5% of tachyzoites by chance form multiple (3-4) daughters per division round: the timing of daughter budding is still synchronized with the other parasites in the vacuole despite the fact that these tachyzoites have undergone two rounds of S/M phase (Hu et al., 2004). For signaling purposes, these paradoxical “multi-daughter endodyogenic” parasites are still in the synchronized cycle of “S/M coupled to budding” state consistent with endodyogeny. Accumulating insights from various mutants displaying increased incidences of “multi-daughter endodyogenic” parasites (Dubey et al., 2017) suggests that such parasites fail to start budding because they are missing membrane building blocks to assemble the daughter IMC, and then slip into the next cell cycle while remaining in the “S/M coupled to budding” state.

When karyokinesis does not follow each S/M phase polyploid nuclei are formed. In *S. neurona* endopolygeny, karyokinesis does not occur at all till the onset of budding, but in Heamosporidioan sporogenesis, karyokinesis is more optional, which leads to a mix of nuclei with various levels of ploidy. However, the mitotic cycle within each nucleoplasm appears to be coordinated (Gerald et al., 2011; Roques et al., 2015), suggesting this level of control is not at the centrosome level, but likely the chromatin or nucleoplasm level. The control and mechanism of karyokinesis are not understood, which is also the case in well-studied model eukaryotes dividing by closed mitosis such as *Aspergillus nidulans*, fission yeast, and baker’s yeast. The challenge in polyploid nuclei is to keep the multiple sets of chromosomes together so they can be accurately partitioned into the daughters. The solution is that the chromosomes remain clustered throughout the cell cycle by tethering the centromeres to the nuclear lamina. For instance, in *T. gondii* the 13 centromeres and associated kinetochores remain clustered at the centrocone, a nuclear envelope fold that houses the spindle microtubules during mitosis (Brooks et al., 2011; Farrell and Gubbels, 2014). Similar observations have been made throughout the *Plasmodium* life cycle using kinetochore markers (Pandey et al., 2019). Strayed chromosomes are rarely observed but individual centromeres are seen during interphase in the *Plasmodium* schizogony in the red blood cell. However, all centromeres cluster together again at the spindle pole before entering mitosis (Hoeijmakers et al., 2012). Clearly, centromere clustering and sequestration at the nuclear envelope through the kinetochores is an effective strategy to warrant that complete sets of chromosomes are maintained in polyploid nuclei.

None of the above addresses why there are two different forms of budding; i.e. internal vs external. We will try to address this by looking at the phylogeny. The first question is whether either internal budding or external budding are an innovation or ancestral process. Since not enough cell division details are known for Apicomplexa outside those groups included in Fig 1, a firm answer is not possible. In favor of loss of internal budding is the generally more reduced genome and streamlined biology found in the *Plasmodium* spp. and the piroplasms (e.g. host cell invasion (Gubbels and Duraisingh, 2012)). Alternatively, in favor of innovation of internal budding in the cyst-forming Coccidia is the putative advantage evidenced during *T. gondii* endodyogeny. In this case, the mother parasite remains invasion- and egress-competent throughout most of the replication cycle, hence increasing the resilience of the parasite in the dynamic host system (i.e. immune attacks) to move between host cells (Gaji et al., 2011). In contrast, during *S. neurona* endopolygeny the micronemes disappear halfway through the division process, which depletes their invasion capacity (Vaishnava et al., 2005); hence, this model does not hold true for the polyploid internal budding modes. Alternatively, it can be argued that the maintenance of a cortical cytoskeleton in the mother makes the parasites more resistant to mechanical stress, but it is not immediately obvious how this sets the tissue cyst-forming Coccidia apart from parasites dividing by external budding. As all of the Coccidia have oral transmission routes, each species exhibits asexual stages in gut epithelial cells; indeed, the non-cyst forming Coccidia complete their whole cycle in the intestinal epithelium. This revelation suggests that the differentiation of internal from external budding might be coupled to the innovation of tissue cyst formation. The requirements for cyst formation comprise dispersion throughout the host beyond the gut-epithelium and assembly of the cyst in neuronal and/or muscular tissue, which do not have obvious links to division modes. Both the tissue cyst inhabiting bradyzoite stages of *T. gondii* and *S. neurona* (pre-bradyzoite metrocytes as well) undergo endodyogeny; only the acute stage of *S. neurona* divides by endopolygeny (Dubey et al., 2001). In addition, although the acute stage of *S. neurona* dissolves the vacuolar compartment and replicates in the host cell cytoplasm, bradyzoite multiplication and cyst formation occur within a vacuole (Jakel et al., 2001). Cyst expansion has to occur gradually so as not to compromise the cyst wall, which obviously is more compatible with endodyogeny than endopolygeny. Notably, bradyzoite replication within *T. gondii* cysts is asynchronous and is consistent with this model (Watts et al., 2015).

An outlier is *Eimeria callospermophili*, a rodent parasite that replicates by endopolygeny rather than schizogony, and as far as known, does not form tissue cysts (Roberts et al., 1970; Hammond, 1973). *E. callospermophili* undergoes several rounds of S-phase that are each followed by karyokinesis to produce 4-10 nuclei per schizont while the mother’s cytoskeleton is maintained. In synchrony, two daughters per nucleus bud internally and the plasma membrane is acquired when the daughter are about 1/3 developed, resulting in 8-20 merozoites. This suggests internal budding was likely present before the advent of tissue cyst formation. However, the phylogenetic position of this parasite is on the edge of the *Eimeria* spp. (Fig 1). To complicate matters further, genome information has demonstrated that the *Eimeria* spp. and the *Isopora* spp. branch within each other and that neither are monophyletic clades (Kvicerova and Hypsa, 2013). Furthermore, the mechanism of cortical budding occurs across the Coccidia sexual cycle during microgametocyte formation (Ferguson et al., 1974; Ferguson et al., 2008). Hence, parasites can exhibit both cell division modes at different stages of their life-cycles – internal and external budding. We also know that sporozoite formation in *T. gondii* occurs through an internal budding mechanism in the absence of a maternal cytoskeleton (Ferguson et al., 1979b; Dubey et al., 2017). Interestingly, the timing of plasma membrane association with the cytoskeleton is much earlier in this situation compared to the other asexual stages occurring in presence of a maternal cytoskeleton, which indicates the timing of events is flexible. Taken together, there appears to be a continuum between the morphogenic features of the Coccidia, suggesting that there is a significant level of plasticity in biology and division modes. Further insights may result from the investigation of parasite species straddling the two division modes; for example, the *Goussia* spp. bridging the cyst-forming and non-cyst forming Coccidia (Fig 1) (Barta et al., 2001). The *Goussia* spp. are found in fish, reptiles and amphibians, which have not been extensively studied at the cell biological level (Rosenthal et al., 2016). Overall, why the two different extant forms of budding exist cannot be answered with satisfaction based on the factors we considered.

This leaves the question of what the mechanistic differences are between internal and external budding. The key question here is how parasites transitioned between the general strategies of external vs internal budding. A principal difference is rooted in the localization of the centrosome. In cortical budding the centrosomes reorient to the plasma membrane, whereas during internal budding they remain in the cytoplasm. A simple model suggests that the presence of the mother’s cytoskeleton physically prevents access of the centrosome to the plasma membrane. Yet, it is unlikely such a simplistic model fully describes this complicated process. A structure anchoring the centrosome to the cortical cytoskeleton has been described for *T. gondii* endodyogeny; a striated fiber assemblin (SFA), extending from the centrosome to the conoid (Francia et al., 2012). SFA fibers are typically found in the flagellar assembly of algae, where they contribute to orientation of the basal body rootlet system relative to other subcellular structures (Francia et al., 2015). During *Eimeria necatrix* schizogony, a dense structure anchors the centrosome to the plasma membrane. This structure migrates basally along with the cortical microtubules during the progression of daughter bud assembly (Dubremetz, 1975), and might be related to the SFA fiber. Such a structure has not been described for *Plasmodium* schizogony in the red blood cell, although the SFA genes are conserved in *Plasmodium* spp. (Lechtreck, 2003). Thus, SFA-like structures anchoring the centrosome to the plasma membrane have been observed in *Eimeria* schizogony (Dubremetz, 1971) and even in *Theileria equi* sporozoite schizogony (Moltmann et al., 1983) and appear to be a differentiating factor between internal and external budding. Further study of SFA genes across the Apicomplexa may reveal the nature of how nuclei are anchored in the buds across the different division modes.

Several more principle difference between internal budding relative to cortical budding exist. For example, the mother and daughter cytoskeletons must be differentially stabilized upon completion of budding. What are the putative mechanisms? It has been shown that a timely regulated proteolysis which removes the C-terminus of the major network component, IMC1, coinciding with conversion of the network from a detergent-labile to a detergent-resistant state late in *T. gondii* daughter cell development (Mann et al., 2002). On the other hand, ubiquitination of the mother’s cytoskeleton marks it for destruction (Silmon de Monerri et al., 2015; Dhara and Sinai, 2016). Although the former informs about maturation and the latter about destabilization, neither informs directly on the basis of differential stability. In parallel, some of the maternal building blocks (e.g. IMC proteins, glideosome) are recycled in the final growth spurt of the daughters (Ouologuem and Roos, 2014; Periz et al., 2019). An additional mechanism is provided by differential components present on either mother (e.g. MSC1b (Lorestani et al., 2012), GAP45 (Gaskins et al., 2004), or IMC7, 12, 14, 17, 18 and 20 (Anderson-White et al., 2011; Chen et al., 2015a)) or daughter parasite (e.g. IMC16, 29 (Chen et al., 2015a; Chen et al., 2016)), or swapping places from mother to daughter (e.g. SPM3, (Samad et al., 2015)). Such differential composition of mother and daughters may be factors in differential stability, but to date no single factor provides a satisfactory explanation.

Another potential problem posed by the mother’s cytoskeleton during internal budding occurs in the growth phase, where it restricts access to nutrients and complicates the ability to expel waste. This is most relevant for the expanding cell during endopolygeny with large cytoplasmic mass. Although both the apical and basal extremes of the cytoskeleton have openings toward the plasma membrane, they are relatively small and it is not clear whether they are sufficient for the level of exchange needed. However, it is not clear whether the sutures between the alveoli are permeable for diffusion of small molecules, which would void this argument. Either way, a set of apical annuli largely composed of AAP proteins residing in the IMC sutures were recently suggested to function as pores across the IMC (Engelberg et al., 2019; Lentini et al., 2019). In support of this hypothesis is the narrow conservation of AAP protein orthologs only in the Sarcocystidae, indicating their function is most likely in support of internal budding. However, disruption of apical annuli structure resulted in a minor fitness loss of *T. gondii* tachyzoites, which might be because endodyogeny poses rather limited demands on exchange, especially compared to endopolygenic replication modes where the mother cytoskeleton could pose a much larger barrier. An alternative role for the apical annuli is as a gateway for a burst in dense granule secretion following completion of invasion (Carruthers and Sibley, 1997): electron microscopy studies place dense granule release at this moment in an apical location consistent with the position of the annuli (Dubremetz et al., 1993). Additional studies on the role of the annuli during endopolygeny are needed to firmly differentiate between these two potential roles.

In conclusion, we define two principally different asexual cell division modes, external and internal budding, which can both produce as few as two daughters per division round, but in most situations produce many more daughters per division round. The bipartite centrosome model and the identification of several regulators of cell cycle progression and checkpoints in recent years provide a framework to explain these division modes, however the evolutionary history and cell biological features defining, organizing and executing the various division modes are much less clear. Expanding genetic and cell biological toolboxes for parasites representing the various division modes provide exciting future avenues toward resolving the exotic apicomplexan cell division modes and shedding light on the evolutionary pressures that select for diversification and choices for different division modes at different developmental stages.

## 5 Conflict of Interest Statement

The authors declare that the research was conducted in the absence of any commercial or financial relationships that could be construed as a potential conflict of interest.

## 6 Author Contributions Statement

MG, HW, DH and MD conceived the experiments, HW performed the *C. suis* experiments, SD and DH performed the *S. neurona* experiments, MG, CK, AP, BE, and CB performed the *Babesia* experiments, KE and MG performed SIM microscopy, BE and IC performed electron microscopy, MG wrote the manuscript and all authors edited the manuscript.

## 7 Funding

This study was supported by National Science Foundation (NSF) Major Research Instrumentation grant 1626072, National Institute of Health grants AI110690 (MJG), AI110638 (MJG), AI128136 (MJG), AI144856 (MJG), and AI128480 (MTD), an American Heart Association pre-doctoral fellowship19PRE34380106 (CDK), a Profillinien start-up grant of the University of Veterinary Medicine Vienna PP16110262 (HLW), an Australian NHMRC CJ Martin fellowship (BE), a post-doctoral fellowship grant 17POST33670577 (KE), a Knights Templar Eye Foundation Career Starter Award (KE), and USDA NIFA grant 2009-65109-05918 (DKH). The funders had no role in study design, data collection and analysis, decision to publish, or preparation of the manuscript.

## 8 Acknowledgements

We thank Bret Judson and the Boston College Imaging Core as well as Stephan Handschuh and the Imaging Core of the University of Veterinary Medicine Vienna for infrastructure and support, Drs. Naomi Morrissette, Jaime Tarigo, and Jeff Dvorin for discussion, and Drs. David Allred, Kirk Deitsch and Laura Kirkman for sharing reagents.

## Supplementary Material

**Figure S1.**
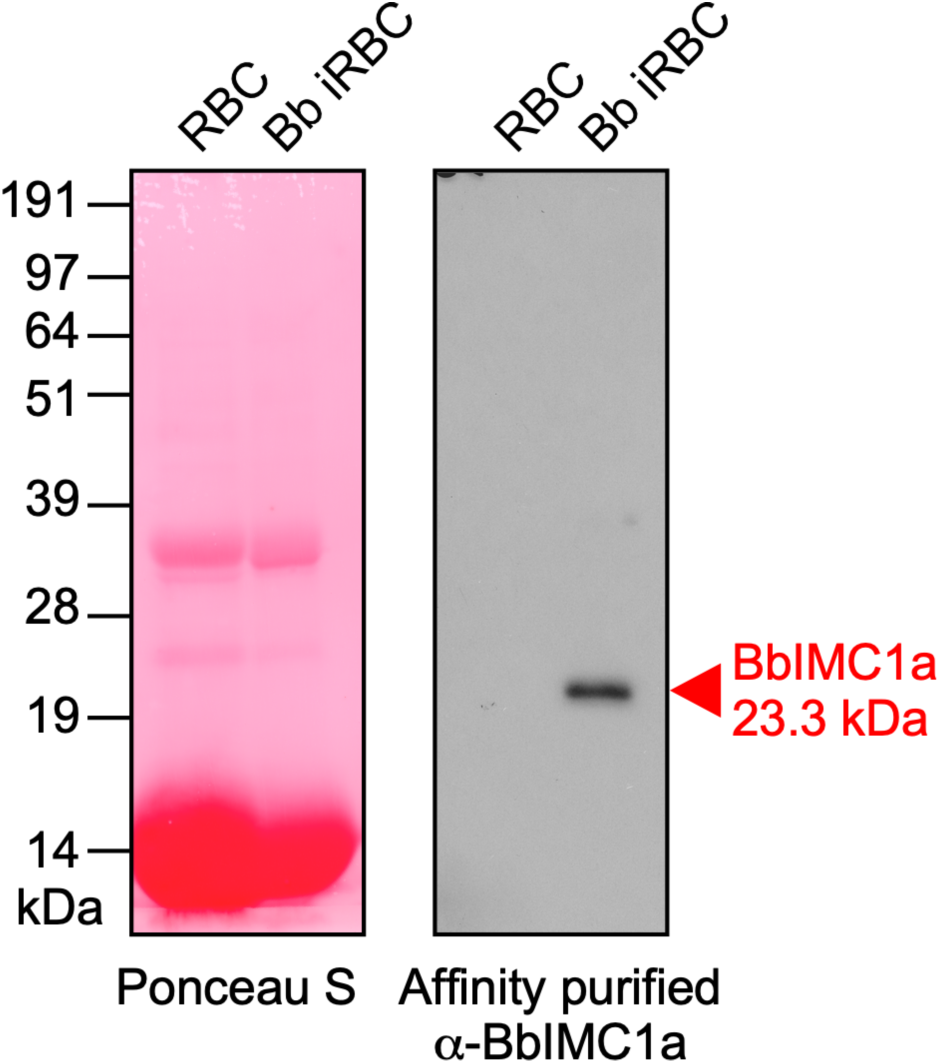
Validation of affinity purified guinea pig antiserum raised against recombinant His6-BbIMC1a by western blot. Left panel: PonceauS staining of the western blot serving as loading control; Right panel: serum affinity purified against recombinant His6-BbIMC1a diluted 1:250. RBC indicates cow red blood cell total lysate; Bb iRBC indicates total lysate of cow red blood cells with a *B. bigemina* parasitemia of 12%. Equal amounts of lysate were loaded across lanes. The predicted MW of BbIMC1a is 23.3 kDa.

## Notes

### Competing Interest Statement

The authors have declared no competing interest.

